# LDEN: Alternating projections for whole genome linkage disequilibrium estimation from next-generation sequencing data

**DOI:** 10.64898/2026.11.16.766929

**Authors:** Syed Muhammad Umer Abdullah, Yanni Sun

**Affiliations:** Department of Electrical Engineering, City University of Hong Kong, Tat Chee Avenue, Kowloon, Hong Kong Special Administrative Region, China

## Abstract

Viral populations often exist as quasispecies of closely-related variants with different mutations. Variants that possess combinations of mutations conferring an overall fitness advantage replicate preferentially. Understanding the true fitness effects of mutations involves disentangling the effects of individual mutations from their genetic background. This problem becomes particularly challenging due to linkage disequilibrium, the non-random association between mutations, which can cause neutral mutations to appear fit and vice versa. Viral populations are typically studied via next-generation sequencing, in which the read length is much shorter than the genome length, rendering long-range linkage disequilibrium unobservable. Estimation of linkage disequilibrium can be performed via population reconstruction, which reconstructs complete sequences before calculating linkage disequilibrium. However, this approach is computationally slow, memory-intensive, and prone to errors, particularly in regions of low coverage. In this work, we present a novel, population reconstruction-independent method for linkage disequilibrium estimation directly from next-generation sequencing data. Our approach uses the observed intra-read linkage disequilibrium values to estimate the unobservable inter-read ones by enforcing mathematical constraints on the linkage disequilibrium matrix. Specifically, because linkage disequilibrium quantifies the covariance between mutant allele frequencies, the linkage disequilibrium matrix must be positive semidefinite. We have formulated linkage disequilibrium estimation as a matrix completion problem and solve it via alternating projections on the constraints of positive semidefiniteness and observed linkage disequilibrium values. This allows the propagation of information from observed intra-read associations to the unobservable inter-read ones without reconstructing the individual variants. We demonstrate that our method yields higher accuracy while being two orders of magnitude faster compared to population reconstruction-based estimation. We demonstrate the performance of the method on both simulated as well as real data showing that it can estimate linkage disequilibrium from a variety of populations.

## 1 Introduction

Viral evolution is characterized by high mutation rates, generally resulting from the lack of high-fidelity proofreading mechanisms capable of correcting mutations during replication. As a consequence, viral populations rapidly acquire mutations and exist as quasispecies of diverse groups of genetically distinct variants competing for the same resources. While most mutations result in nonviable variants, some mutations or combinations of mutations result in a fitness advantage for the variant. Variants with higher fitness advantage survive in the long-term. The identification of variants harboring advantageous mutations, followed by the characterization of those mutations, is a crucial step in understanding viral pathogens. However, the presence of interactions between mutations renders this problem particularly challenging. Mutations can both compete with and enhance each other, meaning that the observed prevalence of an individual mutation reflects not only its inherent fitness but also the genomic background on which it occurs. Neutral or even deleterious mutations may hitchhike alongside beneficial mutations and consequently appear at high prevalence in the population^1^. Conversely, beneficial mutations may be purged from the population due to competition with fitter mutations^2^. When mutations interact with each other in a non-random manner, these are said to be in linkage disequilibrium (LD)^3^. LD is prevalent in nature and has been documented in a number of viral pathogens of clinical interest, such as the human immunodeficiency virus (HIV)^4^, the hepatitis C virus (HCV)^5^, the influenza virus^6,7^, and the severe acute respiratory syndrome coronavirus 2 (SARS-CoV-2)^8^. Accurately estimating LD is therefore essential for understanding the inherent selective nature of individual mutations and for distinguishing truly virulent variants from those that merely ride to prominence due to the presence of linked loci.

Viral populations are typically sequenced via next-generation sequencing (NGS) and numerous viral datasets sequenced using this technology are available in public repositories. NGS however suffers from the technological limitation of read length, which is dictated by the sequencing platform and the specific reagent chemistry employed. The read length of NGS is typically much shorter than viral genomes, for example the 300 base pair (bp) read length of the Illumina MiSeq platform is insufficient to read the entire HIV genome, which is 9719 bp in length, in a single read. Consequently, the short read-length of NGS renders the majority of the LD information across the viral genome unobservable.

Existing methods of LD estimation^9–11^ suffer from poor scalability and cannot be applied even to small viral genomes. To address this limitation of LD estimation methods, we had previously presented MPL-R^8^. MPL-R first subsamples the NGS reads, performs parallelized population reconstruction of subsampled populations, and obtains the LD from the reconstructed sequences, yielding a fast and scalable approach for estimating unobservable inter-read LD values. MPL-R, however, is affected by two notable limitations.

The first limitation is that despite the speedup attained via parallelized population reconstruction, the reconstruction process remains resource intensive. The population reconstruction algorithm in MPL-R, Quasirecomb^12^, requires loading the entire dataset into memory, which poses limitations when handling large datasets. The wall clock time of Quasirecomb also scales quadratically (*O*(*N*^2^)) as the number of unique sequences in the population increases. More fundamentally, while population reconstruction solves the problem of LD estimation, it does so via the estimation of the identity and prevalence value of every sequence in the population, which is not actually needed for this task. LD is a two-point metric that only requires computation of single- and double-mutant allele frequencies of the variable loci. Population reconstruction, by contrast, reconstructs the nucleotide identities and prevalence at all loci of the genome, which not only increases the wall clock time but also generates extraneous data not necessary for the problem at hand. The second limitation of MPL-R is that population reconstruction introduces errors of its own, particularly in genomic regions where the coverage is low. Thus, MPL-R, despite being the fastest method that can estimate unobservable inter-read LD from NGS data, leaves considerable room for improvement.

These limitations can be addressed if LD can be estimated without population reconstruction. To perform reconstruction-free LD estimation, we use the property that both the observable LD values and the unobservable ones are inter-dependent as both form a covariance matrix generated by the underlying viral population. This imposes the mathematical constraint that the complete LD matrix must be positive semidefinite. Thus if we can use the observable LD values to estimate the unobservable ones such that the complete LD matrix is a covariance matrix, we can perform LD estimation without performing population reconstruction.

In this work, we introduce LDEN (whole genome LD estimation from NGS data), a framework that takes NGS reads as input, estimates intra-read LD values from the reads, and then uses these known LD values to estimate the unobservable inter-read LD across the entire genome via convex optimization. LDEN estimates the complete LD matrix by enforcing two constraints, the observable intra-read LD values must be preserved, and the resulting complete LD matrix must be a covariance matrix. This formulation maps the biological problem of estimating unobservable associations between mutations into a matrix completion problem. Unlike population reconstruction, LDEN does not infer individual variants. Instead, it directly estimates the pairwise associations between mutations.

We demonstrate that LDEN accurately estimates LD over a range of simulated viral populations and sequencing conditions. We also demonstrate that LDEN is orders of magnitude faster compared to population reconstruction-based LD estimation and reveals biologically relevant LD patterns in real patient data.

## 2 Materials and methods

This study addresses the problem of estimating LD across the entire genome from NGS data without population reconstruction. The primary idea is that observable intra-read LD values constrain the unobservable inter-read ones which can be estimated by enforcing global constraints on the LD matrix. Specifically, the LD matrix must be positive semidefinite as it measures the covariance between the mutant allele frequencies.

The proposed solution is built upon the alternating projections method (APM)^13^ originally developed in finance to estimate unobservable correlations between stock returns. APM was designed to use partially complete correlation matrices to approximate full correlation matrices that best describe the observed correlations. APM alternatively projects the observed correlation matrix onto two convex constraints, unit diagonal and positive semidefiniteness. The correlation matrix obtained via repeated iterations satisfies both constraints and is the required full correlation matrix.

To apply this optimization framework to the LD estimation problem, the APM formulation needs to be mapped such that it reflects the statistical properties of LD instead of financial correlations. In finance, the input to APM is a symmetric matrix with partially known off-diagonal correlation values. This matrix is repeatedly projected onto the unit diagonal constraint (representing self-correlations) and the positive semidefinite constraint to obtain the nearest correlation matrix. In the LD estimation problem, the LD matrices are covariance matrices with known (intra-read) and unknown (inter-read) off-diagonal values. The unit diagonal constraint in APM is modified so that the projection is on to a matrix with the diagonal equal to the variance. However, since all off-diagonal elements are known within the read length, we modified the unit diagonal constraint with that of a band composed of observed LD values. We retained the positive semidefinite constraint from APM. Alternatively projecting the observed LD matrix onto these constraints results in the nearest covariance matrix to the observed LD matrix which is the required full LD matrix. Thus we formulate the problem as finding an LD matrix that contains the observed LD values for locus pairs within one read-length of each other, and approximations of the unobservable LD values such that the overall matrix is positive semidefinite. Supplementary Table S1 summarizes the comparison between the APM framework and the proposed implementation.

We demonstrate that LDEN not only provides accurate estimates of unobservable LD values but is also orders of magnitude faster than population reconstruction-based estimation. This computational advantage stems from the fact that LDEN operates directly on observable two-point statistics, bypassing the requirement of reconstructing full sequences. We further demonstrate the performance of LDEN on populations with different values of evolutionary parameters as well as under different levels of sequencing noise in the NGS reads. Finally, we demonstrate the practical utility of LDEN in inferring biologically meaningful insights from real data of norovirus infection.

### 2.1 Read alignment and partial LD calculation

NGS reads are aligned to a reference sequence, sorted, and indexed via samtools^14^. To reduce the effect of sampling errors, read coverage i.e. the number of reads covering a locus, is used to filter out poorly covered loci (covered by less than 5 reads). Loci exhibiting low variation (less than 1% change in the single mutant allele frequencies) are also removed from subsequent analysis, as these don’t contribute much to LD estimation and unnecessarily increase the number of computations.

The coefficient of LD, *D*, is calculated as *D* = *f*_*i j*_ *f*_*i*_ − *f* _*j*_. Here *f*_*i*_, *f* _*j*_ are the single mutant allele frequencies at loci *i, j*, and *f*_*i j*_ is the double mutant allele frequency at the locus pair (*i, j*). In NGS data, coverage varies across loci. If *f*_*i*_, *f* _*j*_, and *f*_*i j*_ are calculated over different *n*_*i*_, *n* _*j*_, and *n*_*i j*_ reads covering the respective loci and loci pairs, then the resulting value of *D* may violate the axioms of probability due to inconsistent denominators. To circumvent this issue, we compute *f*_*i*_, *f* _*j*_, and *f*_*i j*_ using only *n*_*i j*_ reads that cover both loci simultaneously, ensuring the denominators are consistent.

### 2.2 Linear shrinkage for low-coverage

When the number of reads covering a pair of loci becomes small, the resulting LD estimates are affected by statistical noise of the order of 1*/n*_*i j*_. This can lead to severe overestimation or underestimation of true LD values. As the coverage of NGS typically follows a trapezoidal profile, with loci close to the ends of the genome having lower coverage values compared to loci close to the center, this problem requires explicit treatment. To minimize the effects of low coverage, we employ a shrinkage estimator on the frequencies as

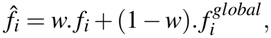

and

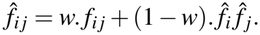

Here 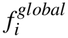 represents the single mutant allele frequency obtained via all reads that cover locus *i, w* is a pairwise coverage-dependent shrinkage term, and 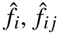, are, respectively, the shrunken estimates of the single and double mutant allele frequencies used in subsequent analysis. Intuitively, *w* quantifies how informative *f*_*i*_ and *f*_*i j*_ are. For high values of pairwise coverage, *w* approaches 1, and 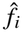 becomes close to *f*_*i*_, whereas for low pairwise coverage, 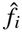 approximates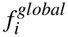. Similarly, for high values of pairwise coverage 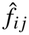 is essentially taken as *f*_*i j*_, whereas when the pairwise coverage is low, *f*_*i j*_ is assigned a lower weight and 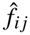 is shrunk towards the product 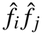 (i.e. towards linkage equilibrium) to avoid sampling errors. The shrinkage weight *w* is defined as *w* = *n*_*i j*_*/*(*n*_*i j*_ + *λ*), where *λ* is a user-defined shrinkage factor. In our tests, we set *λ* empirically based on the noise levels in the sequencing data.

### 2.3 Alternating projections method

The partial LD matrix obtained directly from the reads does not satisfy the positive semidefinite property required for covariance matrices, and hence, is not a valid covariance matrix. However, it can be used to obtain a covariance matrix that best explains the observed LD values while satisfying the positive semidefinite constraint. Formally, for the observed LD matrix *D*, and a candidate covariance matrix *C*, the problem is to minimize the distance *γ*(*D*) between *D* and *C* such that

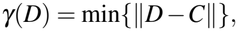

where the norm represents the squared Frobenius norm, i.e.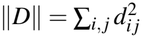. The matrix that achieves this minimum distance is the required covariance matrix. Such a matrix is unique and its existence is a known result in convex optimization^15^. The matrix can be obtained by exploiting two properties, i.e. the LD matrix must positive semidefinite and its entries within the read length must equal the observed LD values, mathematically

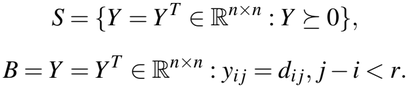

Here *r* represents the read length, *n* is the total number of loci, and *Y* ⪰ 0 indicates that *Y* is positive semidefinite. Both *S* (the positve semidefinite constraint) and *B* (intra-read entries equal to the observed entries constraint) are closed convex sets, therefore their intersection is also a closed convex set. The matrix that satisfies both constraints can be obtained iteratively by alternately projecting *D* onto *S* and *B* (Fig. 1).

**Figure 1.**
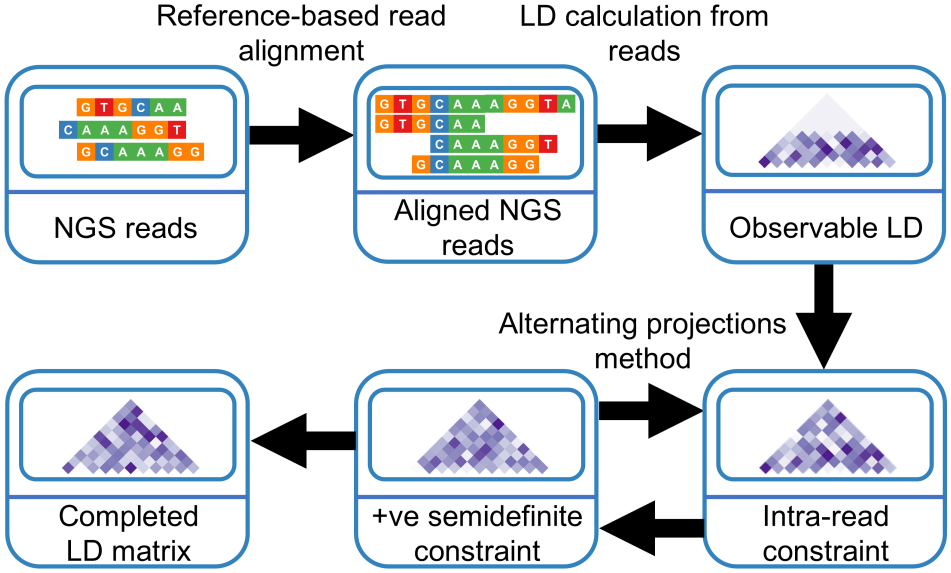
LDEN processing schematic. NGS reads are aligned to a reference sequence, sorted, and indexed. Observable LD values are calculated between loci pairs jointly covered by individual reads and the LD matrix is partially populated with these values. The entries corresponding to the inter-read LD values are missing in the LD matrix. LDEN completes the LD matrix by alternating projections on two constraints, preservation of the observed LD entries and positive semidefiniteness of the complete LD matrix.

In the alternating projection method, an eigendecomposition of *D* is first obtained and all negative eigenvalues are set to a small positive value to enforce the positive semidefinite constraint. This is followed by replacing the observed LD entries, corresponding to intra-read loci pairs, back into the iterate, thereby enforcing the intra-read constraint. These two projections are alternated until the difference between the Frobenius norm of successive iterates falls below a specified convergence threshold, *ε*, or the maximum number of iterations, *M*, is reached. Empirical tests on convergence discuss these parameters in detail in the Results section. The complete algorithm is presented below.

#### Algorithm 1

The LDEN algorithm

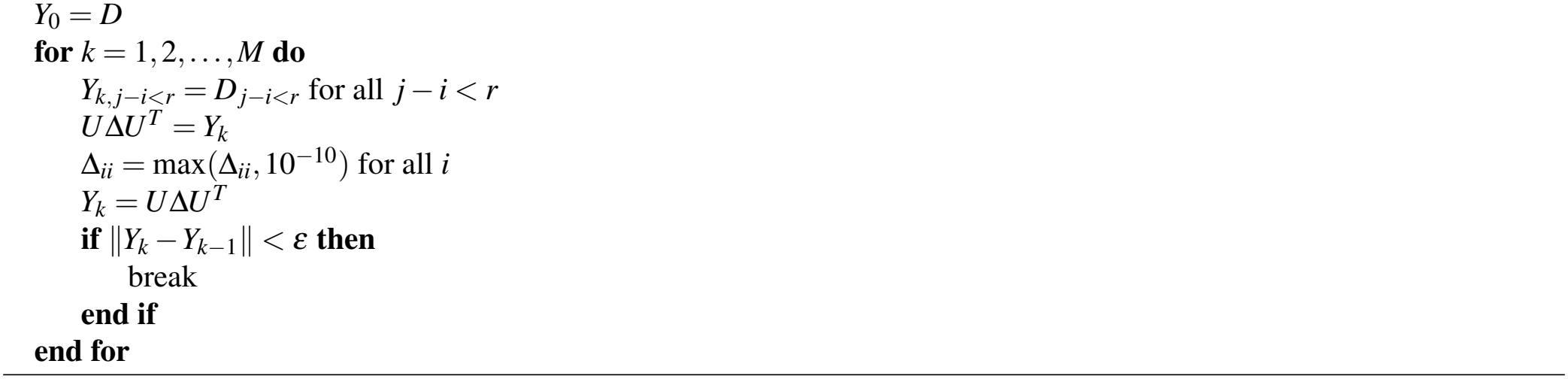

Upon convergence, the final iterate *Y*_*k*_ is the required completed LD matrix *C*. It is positive semidefinite, has all observed intra-read LD values, and has the remaining entries consistent with the defined constraints. Theoretically, iteration should continue till *C* = *Y*_*k*_ satisfies *γ*(*D*) = min {∥*D* − *C*∥}, but we will demonstrate later that this requirement can be relaxed, and iteration can stop before without significant loss in accuracy.

### 2.4 LD estimation from paired-end reads

In addition to single-end sequencing, many NGS sequencing platforms support paired-end sequencing, in which both ends of a genomic fragment are sequenced. Paired-end sequencing provides two key advantages compared to single-end sequencing: (i) it allows the observation of the same genomic fragment from both directions, which provides more information about the composition of the fragment; and (ii) paired-end reads capture LD across longer ranges compared to single-end reads, as the two reads in a pair originate from the same fragment and can span regions that cannot be covered by single-end reads. Consequently, incorporating information from paired-end reads into the LDEN framework is expected to improve the accuracy of the recovered LD matrix by increasing the proportion of the directly observable LD entries. We incorporated the functionality of utilizing fragment-level LD observed by read pairs into the LDEN framework. Instead of calculating LD over individual reads, LDEN can also calculate it over read fragments composed of paired reads. This approach extends the range of the observable LD beyond the length of a single read, increasing the genomic distance across which LD can be directly observed. Similar to single-end reads, paired-end reads are affected by sampling noise, particularly when the number of read pairs covering a locus pair is small. We therefore employ a pairwise coverage threshold, considering only those locus pairs that were covered by at least five read pairs. Locus pairs covered by fewer than five pairs are excluded from the observed set of loci and are left for imputation by the alternating projections algorithm. Additionally, we apply the same linear shrinkage estimator described in Section 2.2 to the double mutant allele frequencies and use the same value of the shrinkage factor as the singe-end reads case. This ensures consistent regularization in the two-step approach and uniformly mitigates the effects of statistical noise in regions with low coverage.

### 2.5 Simulated data

#### 2.5.1 Overview

We simulated data based on realistic viral populations to perform analytical validation of LDEN. A total of 6 primary datasets were generated, each with varying population parameters. The total number of sequences in every dataset was set to 1000.

#### 2.5.2 Variation in population diversity

For datasets 1 through 4, a random founder sequence of length 500 bp was generated by appending the nucleotides A, C, G, and T, uniformly at random, with each nucleotide having equal probability of occurrence, i.e. 0.25. From this founder sequence, 3 variant sequences were generated by randomly mutating 1%, 2%, and 5% of the loci, respectively. Co-infection dynamics were simulated by assigning prevalence values to each variant according to the distributions shown in Supplementary Table S2. In dataset 1, a single dominant variant (founder 1) comprises the majority of the population, with the remaining prevalence distributed among the other variants. Dataset 2 represents a population with two co-dominant founders. Dataset 3 contains three founders with equal prevalence, whereas dataset 4 comprises four founders with similar prevalence.

#### 2.5.3 Variation in genome length

In datasets 5 and 6, the prevalence values were kept the same as those of dataset 1, while the genome length was increased to 1000 bp and 1500 bp respectively. These datasets were generated to evaluate the performance of LDEN as genome size was increased.

#### 2.5.4 Wright-Fisher time-series simulation

We also simulated a time-series dataset via a Wright-Fisher evolutionary model^16,17^. A population of 1000 sequences, each of genome length 500 bp was considered with 460 neutral loci under zero selection, and 20/20 randomly distributed loci with beneficial/deleterious mutations of selection strength ± 0.025 respectively. Mutations were simulated with a probability of 10^−4^ mutations per locus per generation. The population was allowed to evolve for 400 generations and sampling was performed at every 50 generations, yielding a total of 9 observed generations.

#### 2.5.5 NGS reads simulation

NGS reads were simulated via the ART simulator^18^. Single-end reads of length 150 bp were generated using the error profile of the Illumina HiSeq 2500 platform. For datasets 1 to 4, the reads-per-sequence parameter was set to 5 to generate 5000 reads with a coverage of 1500X. For datasets 5 and 6, the reads-per-sequence parameter was set to 8 and 10 respectively, and 500 reads were randomly removed from dataset 5 via custom code to achieve the same coverage of 1500X as datasets 1 to 4. For the time-series dataset, the reads-per-sequence parameter was set to 15 to obtain 15,000 reads with a coverage of 4500X. We also simulated paired-end NGS reads of 150 bp for dataset 1 by setting the mean fragment length to 200 ± 50. The reads-per-sequence parameter was set to 5 to obtain 5000 reads and custom code was used to select 2500 read pairs from the generated reads to get the same coverage of 1500X as the single-end reads dataset.

#### 2.5.6 Sequencing parameter sensitivity analysis

To investigate the sensitivity of LDEN to sequencing parameters, we generated additional datasets based on dataset 1 with varying read length and coverage values. Specifically, two datasets were generated with read lengths 50 bp and 100 bp, while keeping the coverage fixed at 1500X. Three datasets were generated with coverage values 50X, 500X, and 4500X, keeping the read length fixed at 150 bp.

#### 2.5.7 Statistical confidence

For each dataset and experimental condition, 100 independent Monte Carlo runs were simulated. This replication was done to establish statistical confidence in the results.

### 2.6 Norovirus data

To perform biological concordance analysis of LDEN on real patient data, we performed LD estimation on the data of an immunocompromised patient with GII.4 norovirus infection^19^. The patient, labeled P1 in the original study, had been infected with the GII.4 Sydney[P16] 2012 HuNoV strain of the virus and had been undergoing treatment with favipiravir. The dataset comprises four time-series samples obtained prior to the administration of favipiravir, 10 samples obtained during treatment with 200 milligram (mg) favipiravir, and five samples obtained during treatment with 800mg favipiravir. Sequencing demonstrated that the virus developed multiple mutations during treatment. We chose the time-series samples obtained during treatment with 800mg favipiravir as those contained multiple co-occurring mutations, indicative of LD (Fig. 3 in Kreins *et al*. 2024^19^). These samples were obtained between 433 and 611 days since the first positive sample. Sequencing was performed via the Illumina platform to obtain 151 bp paired-end reads. We chose to analyze the major capsid protein (VP1) as it is known to contain multiple experimentally-determined epitopes. The genomics coordinates (5085-6707) of VP1 were obtained from the GII.4 Sydney[P16] 2012 HuNoV reference sequence (GenBank accession no. JX459908.1). The length of the gene (1623 bp) was much longer than the read length (151 bp) and significant LD information was unobservable. Burrows-Wheeler Aligner-Maximal Exact Matches (BWA-MEM)^20^ was first used to align the reads against the region of the reference sequence corresponding to VP1. The aligned reads were converted to BAM format and sorted via samtools. The mean number of mapped and paired reads per sample was 305540 ±164331, which yielded an average coverage of 28427X±15289X. The mpileup command of samtools and VarScan^21^ were used to obtain the single mutant allele frequencies, from which the variable loci were identified based on the 1% threshold described before. LD estimation was performed via LDEN across all variable loci in the VP1 region. To establish concordance of the estimated LD with biological knowledge, we used the rank ordering of the selection coefficients of the mutations. As nonsynonymous mutations within epitopes are likely driven by immune pressure, correct estimation of their selection coefficients is expected to show a higher number of these in the most strongly selected mutations as compared to random chance. To identify nonsynonymous mutations, we used LoFreq^22^ to obtain the identity of the WT and mutant allele at each locus, followed by custom code to mark synonymous and nonsynonymous mutations. To obtain the coordinates of the epitopes in VP1, we utilized Fig. 1 in Tohma *et al*. 2025^23^. We then used the LD estimated via LDEN and performed linkage-aware selection coefficient estimation of all mutations via marginal path likelihood (MPL)^4^. As a baseline, we also performed linkage-agnostic estimation (LD-agnostic) of the selection coefficients, by setting the off-diagonal entries of the LD matrices to zero, to quantify the improvement achieved by performing LD estimation via LDEN. For selection coefficient estimation via MPL, a bi-allelic model of WT/mutant was adopted. The regularization parameter for MPL was set to 10 following our previous results on this parameter (Fig. S2 in Abdullah *et. al* 2026^8^). The mutation probability from wild type (WT) to mutant was set as 7.4 × 10^−6^ mutations per locus per day based on Matsushima *et al*. 2019^24^, whereas three times this number was chosen as the mutation probability from mutant to WT to account for mutations from three different nucleotides to the WT. The rank ordering of the estimated selection coefficients was verified for statistical confidence via fold enrichment, defined 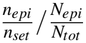, where *N*_*tot*_ is the total number of mutations, *N*_*epi*_ is the total number of epitope-associated nonsynonymous mutations, *n*_*set*_ is the number of the strongest *x*% mutations, and *n*_*epi*_ is the number of epitope-associated nonsynonymous mutations in the strongest *x*% mutations. A fold enrichment value close to 1 indicates random chance, whereas higher values indicate greater likelihood of statistically significant results, quantified by the *P*-value obtained via one-sided Fisher’s exact test. To account for variation in sampling, we also obtained the standard deviation in the fold enrichment by performing non-parametric bootstrap resampling over 1000 bootstrap samples.

## 3. Results

### 3.1 Overview

We evaluated the ability of LDEN in estimating LD from both simulated and real viral sequencing data. Simulated population data was used to evaluate whether sequencing errors and non-uniform coverage affected the estimation of intra-read LD from the reads (Supplementary Section S1.1). We observed that filtering out loci based on observed variation and performing the proposed linear shrinkage improved the estimated LD (Supplementary Fig. S1). We tested the convergence of the algorithm and observed that convergence was fast even when the maximum number of iterations was increased by three orders of magnitude (Supplementary Section S1.2). We also observed that increasing the number of iterations beyond 1000 did not significantly improve the accuracy of the estimated LD and chose to use 1000 iterations as the stopping criterion (Supplementary Fig. S2) in subsequent tests. Change in performance of the LDEN was measured as population parameters were varied. Genetic diversity, controlled by varying the number of founder sequences, and genome length were varied and LDEN was used to estimate LD. Tests were carried out on simulated reads to check for variation in performance as the sequencing quality degraded, specifically, by varying read length and sequence coverage. We then compared the performance of LDEN against population reconstruction-based inference on time-series evolutionary data. Finally, we demonstrated the utility of LDEN on a real dataset of norovirus evolution from an immunocompromised patient.

### 3.2 LDEN recovers valid LD matrices from NGS data

The banded LD matrices directly obtained from NGS reads are not true covariance matrices as these do not satisfy the positive semidefinite constraint required for such matrices. Furthermore, these matrices may also be mathematically inconsistent because replacing unobserved LD values with zeros can violate the theoretical bounds dictated by probability theory. A comparison between the eigenvalues of the banded LD matrices obtained directly from the reads and those of the LD matrices reconstructed via LDEN is shown in Fig. 2. It can be observed that the banded LD matrices exhibit negative eigenvalues, which are impossible for a covariance matrix. On average, around 33.79 ± 3.58% of eigenvalues were negative across all Monte Carlo runs. In contrast, all eigenvalues of the LD matrices recovered via LDEN are nonnegative, satisfying the positive semidefinite property. Thus, LDEN successfully transforms the banded, potentially inconsistent LD matrices observed from NGS reads into valid covariance matrices suitable for downstream evolutionary analysis.

**Figure 2.**
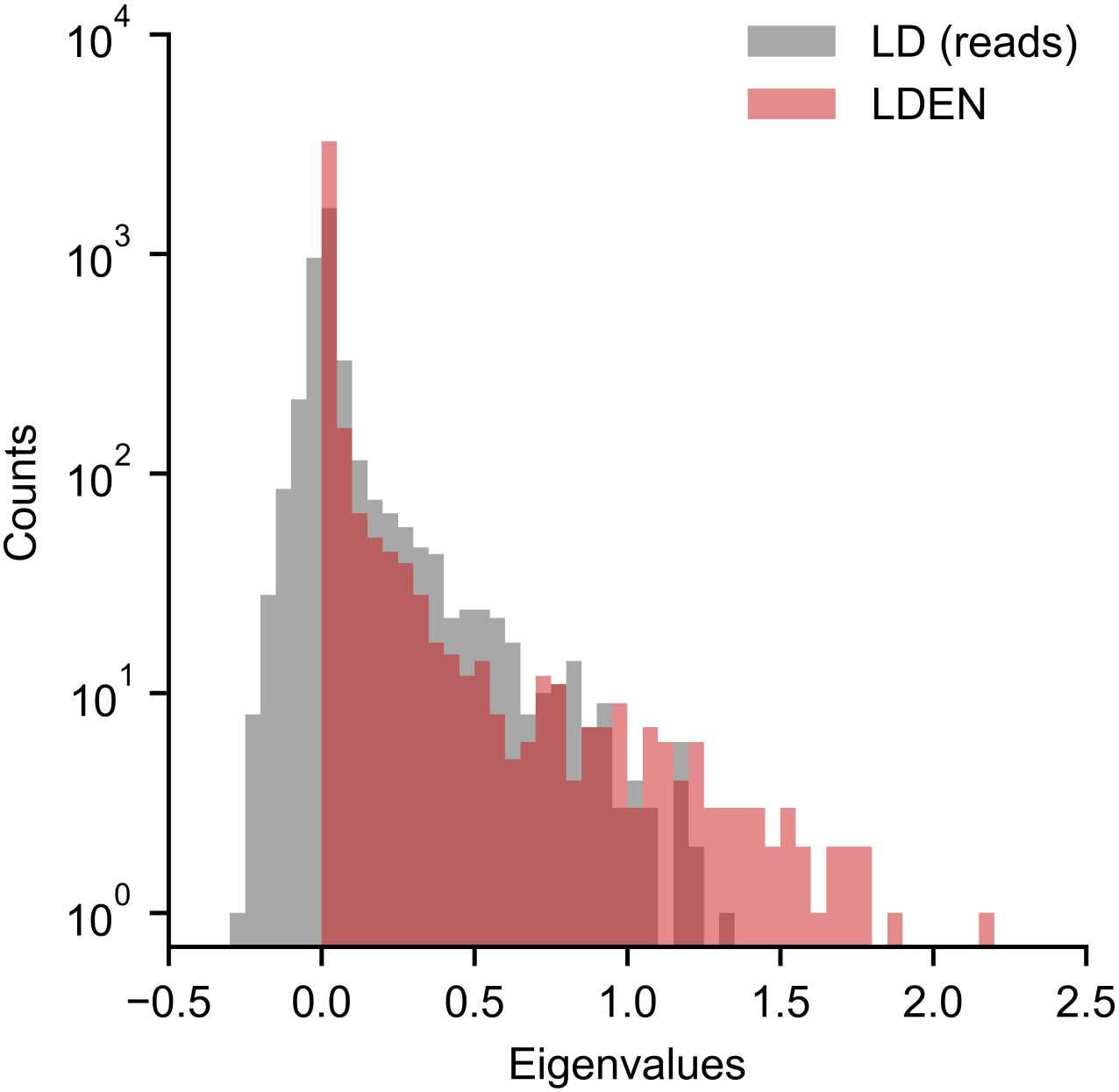
LD matrices estimated by LDEN are positive semidefinite. Histograms showing the eigenvalues of the banded LD matrix obtained directly from the NGS reads (gray) and the LD matrix estimated by LDEN (red). The banded LD matrix contains negative eigenvalues and is not positive semidefinite, whereas the LD matrix estimated by LDEN has all nonnegative eigenvalues. Results are shown for 100 Monte Carlo runs. Each Monte Carlo run consisted of a population of 1000 sequences derived from 4 variants. One founder of length 500 bp sequence was generated randomly while 3 variants were generated from the founder sequence by randomly mutating 1%, 2%, and 5% of the loci, respectively. Prevalence values of each variant follow dataset 1 in Supplementary Table S2. Analysis was performed on single-end NGS reads of 150 bp length with a coverage of 1500X.

### 3.3 The performance of LDEN is correlated with the accuracy of the banded LD information

LD depends on the level of genetic variation in the population, which in turn is influenced by the number of infecting strains, the rates of mutation, recombination, and reassortment of the virus, as well as the time elapsed since infection. LD can also exist over large distances due to three-dimensional interactions between different nucleotides in the viral genome. In addition to the parameters of the population, LD estimation depends on how well the sequencing reads sample the population. Reads may have different levels of coverage which can introduce various degrees of statistical noise into the data. Read lengths also vary across sequencing platforms and determine the extent of LD information individual reads can capture. We evaluated the robustness of LDEN to varying population and read parameters and established benchmarks for the performance of the method. LDEN was observed to be robust to the level of variation in the data (Fig. 3A), however its performance degraded as the genome length was increased while keeping the read length constant (Fig. 3B). The performance of LDEN improved as coverage (Fig. 3C) or read length (Fig. 3D) was increased. This trend in performance can be understood by the fact that higher coverage reduces statistical noise in the observable LD whereas the longer reads enable the observation of more LD information. To explain how longer genome length and shorter read length render more LD information unobservable, we refer to Supplementary Fig. S3. When the read length is fixed and the genome length is increased, the number of unobservable LD values increases as indicated by the bulk of the histograms appearing towards the right of the vertical line (Supplementary Fig. S3A). Conversely, when the genome length is fixed and the read length is decreased, more of the distribution lies towards the right side of the respective vertical line, meaning a greater proportion of LD information becomes unobservable (Supplementary Fig. S3B). Overall, LDEN performs well as long as the NGS reads capture sufficient information from the data. When the genome length is small relative to the read length, or when the read length or coverage is increased, the quality of the information observed by the reads improves, which in turn enhances the accuracy of the LD estimation. These results may be used as guidelines for designing future NGS experiments.

**Figure 3.**
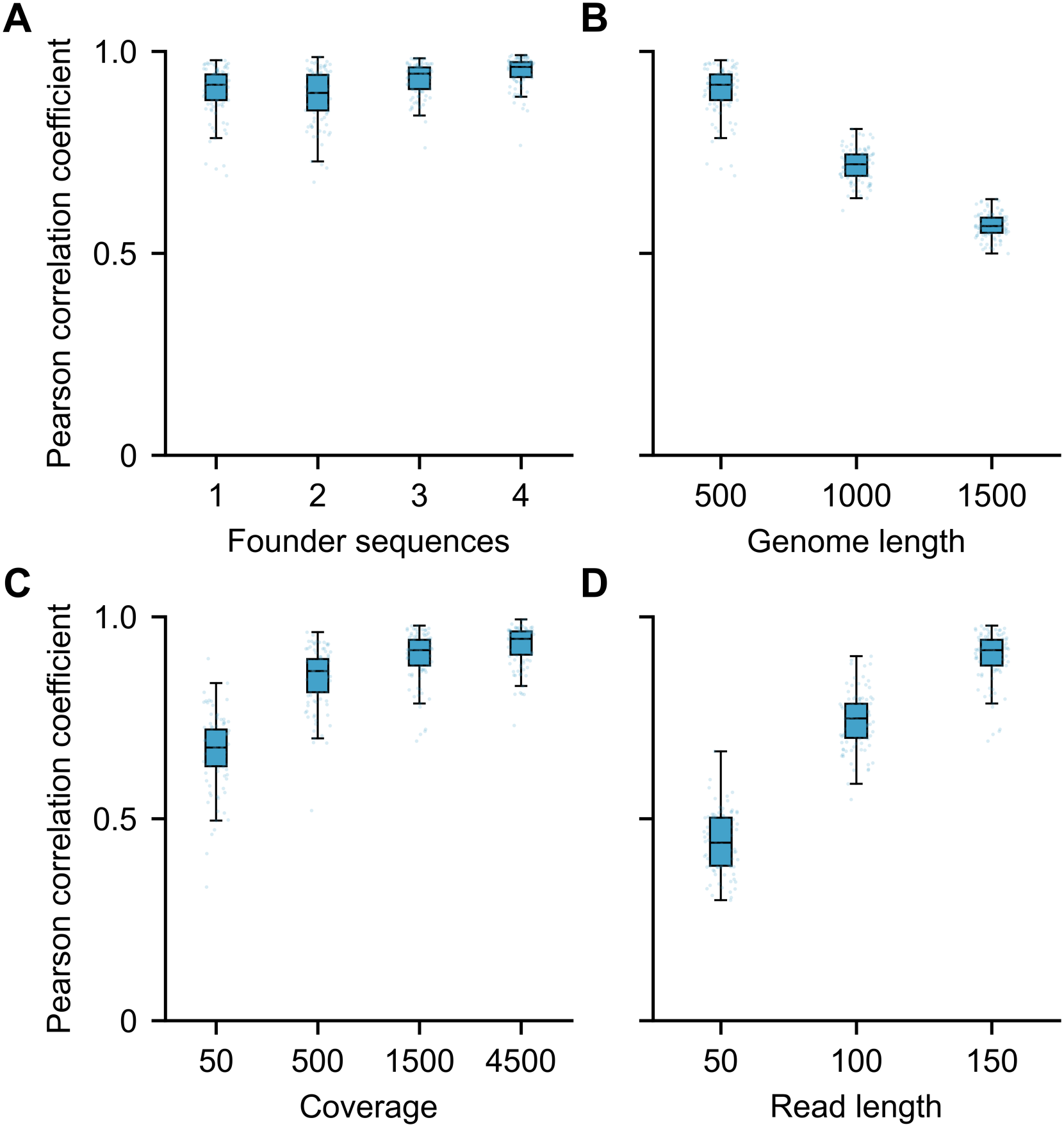
The performance of LDEN is correlated with the accuracy of the information captured by the reads. **(A)** Performance remains similar for different levels of variation in the data. **(B)** Performance degrades as the genome length is increased for the same read length. **(C)** Increase in coverage improves performance. **(D)** Increasing read length improves performance. Results are shown for 100 Monte Carlo runs. Each Monte Carlo run consisted of a population of 1000 sequences derived from 4 variants. One founder sequence was generated randomly while 3 variants were generated from the founder sequence by randomly mutating 1%, 2%, and 5% of the loci, respectively. Prevalence values of each variant are listed in Supplementary Table S2. Analyses on coverage and read length were performed on dataset 1. Analyses on founder sequences and genome length were performed on simulated single-end NGS reads of 150 bp length with a coverage of 1500X.

### 3.4 LDEN outperforms population-reconstruction based estimation

In our previous work on LD estimation, we demonstrated that population reconstruction struggles to accurately estimate LD when genetic diversity is low (Fig. S19 in Abdullah *et. al* 2026^8^). To compare the performance of MPL-R with LDEN, we applied both methods on time-series NGS data. LDEN consistently outperformed MPL-R in terms of the accuracy of the estimated LD (Fig. 4A). In addition to superior accuracy, LDEN completed the entire processing pipeline, from NGS reads to LD estimation, in a median time of around 10 second, which was 2 orders of magnitude faster compared to MPL-R (Fig. 4B). The improved accuracy and computational efficiency of LDEN can be explained by its fundamentally different approach to LD estimation as compared to MPL-R. LDEN bypasses population reconstruction entirely so is not affected by population reconstruction errors. LDEN is faster because it estimates LD directly from the NGS data, whereas MPL-R first performs population reconstruction, which is a computationally intensive process, and subsequently estimates LD from the reconstructed population, which requires much longer wall clock time compared to LDEN.

**Figure 4.**
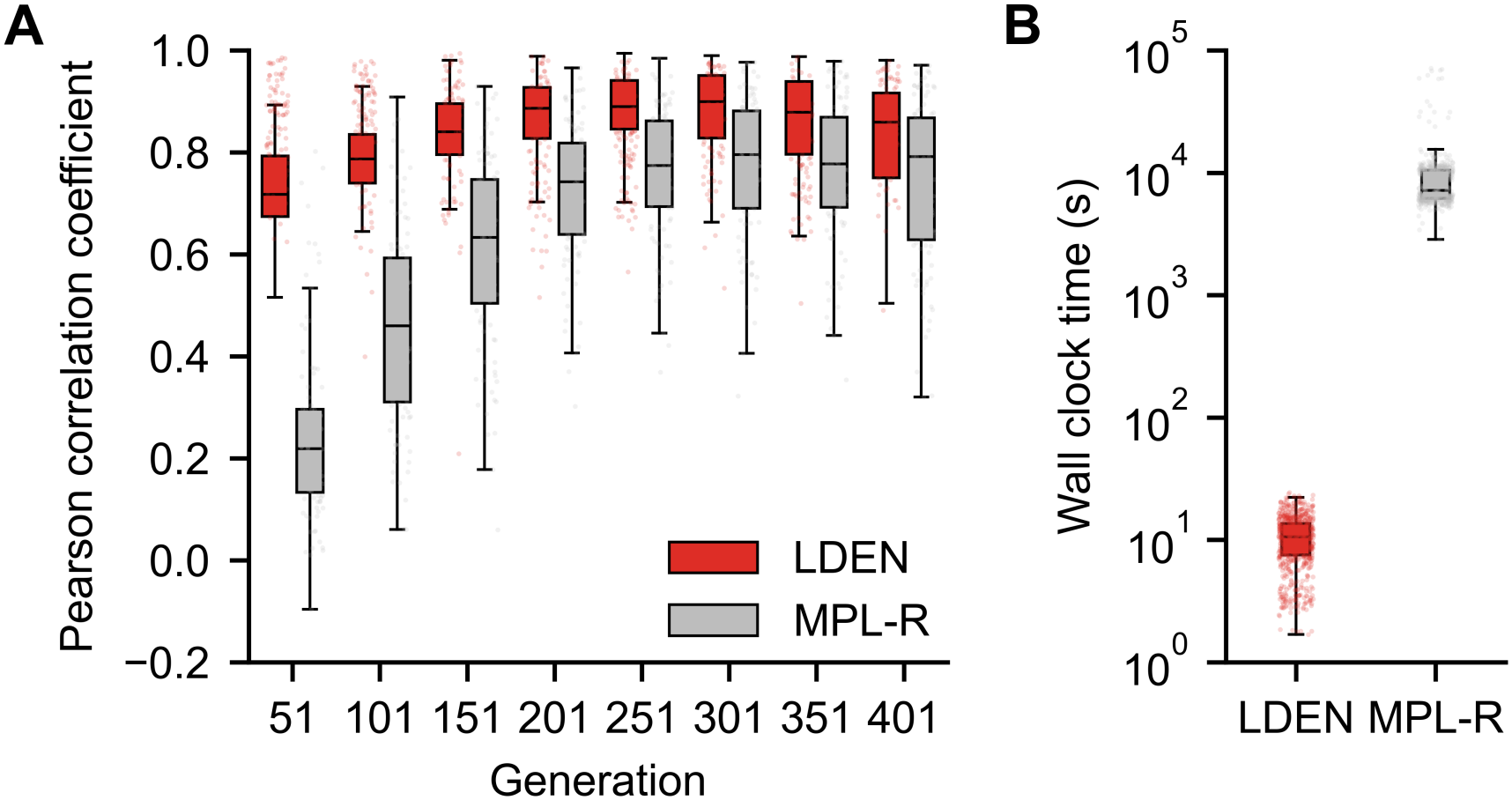
LDEN outperforms MPL-R in both estimation accuracy and computational speed. **(A)** LDEN achieves higher accuracy in LD estimation compared to MPL-R across time-series data. **(B)** LDEN completes the entire processing pipeline two orders of magnitude faster than MPL-R. Results are shown for 100 Monte Carlo runs. Each Monte Carlo run consisted of a population of 1000 sequences evolved from a single randomly generated founder genome of length 500 bp. Mutation was simulated at a probability of 10^−4^ mutations per locus per generation. 460 loci were neutral and 20/20 were under beneficial/deleterious selection with selection coefficients ±0.025. Single-end NGS reads of 150 bp length with a coverage of 4500X were simulated for each time-point. MPL-R was run on 3 bootstrap samples each of coverage 1500X and the results were aggregated.

### 3.5 Paired-end reads improve the accuracy of LDEN

We compared the performance of LDEN on single-end and paired-end reads to assess whether the incorporation of fragment-level linkage information improves the accuracy of LD estimation (Fig. 5). The median Pearson correlation coefficient between the LD matrix recovered from paired-end reads and the ground truth was higher than that between the LD matrix obtained via single-end reads and the ground truth (Fig. 5A). This improvement indicates the greater amount of directly observable LD information captured by paired-end reads, which extends the observable genomic distance beyond the length of individual reads. In terms of computational cost, LD estimation using paired-end reads took a similar time as the single-end case, which is expected due to the same number of reads in both sets. The median wall clock time required to process paired-end reads was approximately 0.7 second per Monte Carlo run (Fig. 5B), which is negligible compared to performing population reconstruction. These results demonstrate that paired-end reads offer improved accuracy of LD recovery without increasing the wall clock time.

**Figure 5.**
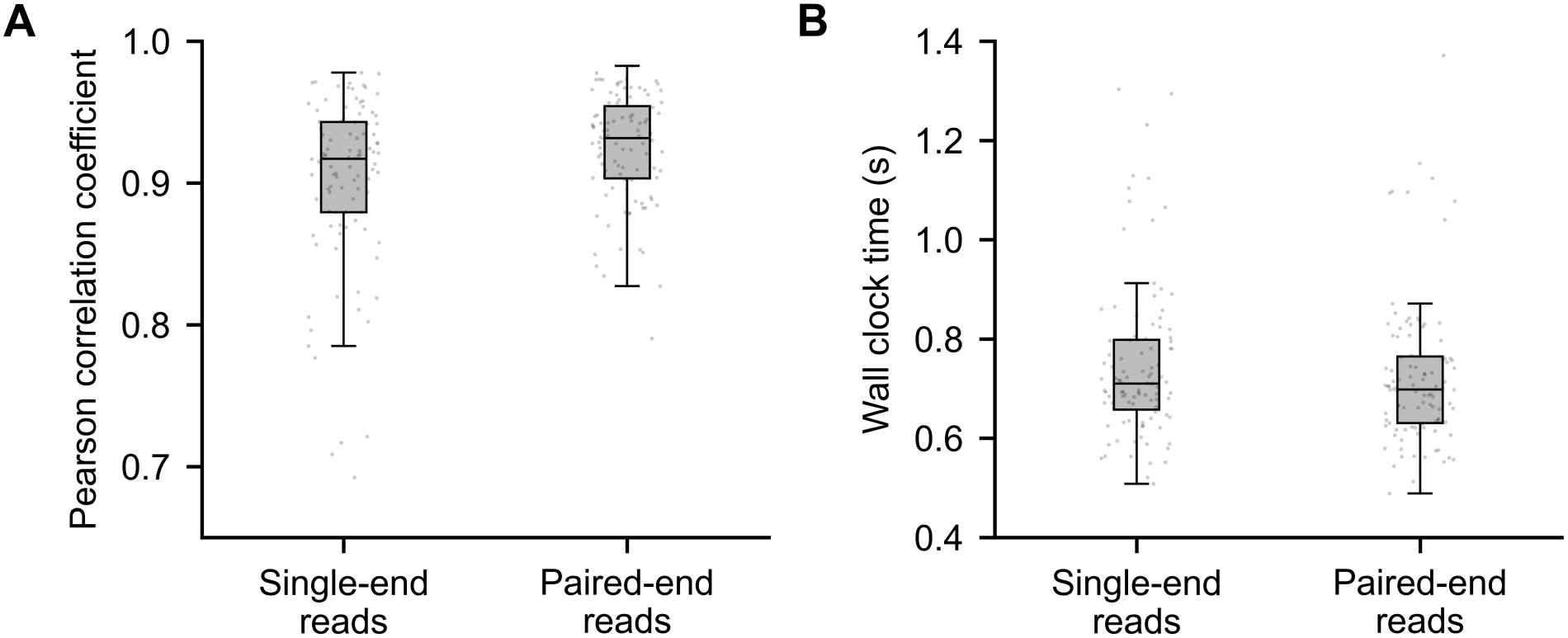
LDEN yields more accurate results using paired-end reads. **(A)** The accuracy of LD estimation is higher when using paired-end reads as compared to single-end reads. **(B)** The wall clock time is higher when using paired-end reads but the estimation still completes in less than 5 seconds. Results are shown for 100 Monte Carlo runs. Each Monte Carlo run consisted of a population of 1000 sequences derived from 4 variants. One founder sequence was generated randomly while 3 variants were generated from the founder sequence by randomly mutating 1%, 2%, and 5% of the loci, respectively. Prevalence values of each variant are listed in Supplementary Table S2. Genome lengths of the sequences were 500, 1000, 1500. Both single- and paired-end reads had a length of 150 bp length with a coverage of 1500X. The mean fragment length for paired-end reads was 200 ± 50.

### 3.6 LDEN reveals known biological insights from patient data

We used LDEN to estimate unobservable LD and used MPL to obtain the selection coefficients for all mutations. We also obtained selection coefficients via LD-agnostic. We then sorted the selection coefficients in descending order based on magnitude. We computed the fold enrichment for both approaches in 1 to 10% of the top strongest selection coefficients. Fig. 6 demonstrates that LDEN consistently yields greater fold enrichment compared to random chance and the estimates are more enriched compared to LD-agnostic in the top 1 to 4% strongest selection coefficients. The highest value of fold enrichment reported by LDEN was 8.12 (Odds ratio=8.95, *P*-value= 2.09× 10^−3^), which was much higher compared to random chance. This corresponded to four epitope-associated nonsynonymous mutations in the top 32 mutations. The maximum value of fold enrichment reported by LD-agnostic was 2.76 (Odds ratio=3.00, *P*-value= 0.16), which was not statistically significant. This corresponded to two epitope-associated nonsynonymous mutations in the top 47 strongest mutations. Overall, the greater number of epitope-associated nonsynonymous mutations identified by LDEN provides biological validation of the accuracy of the recovered LD.

**Figure 6.**
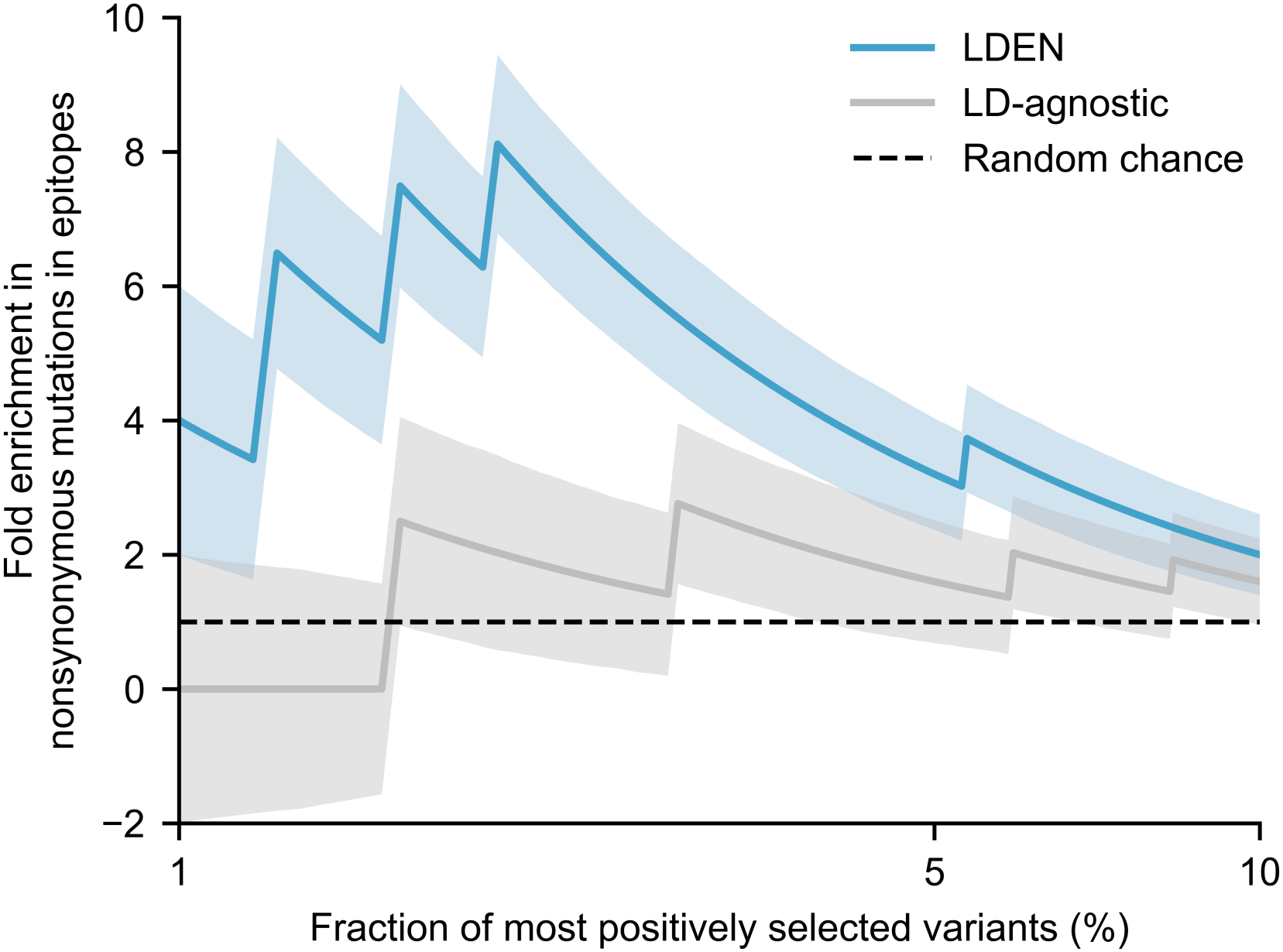
LDEN identifies strong selection on epitope-associated nonsynonymous mutations. Linkage-aware estimation of selection coefficients using the LD estimated via LDEN demonstrates greater fold enrichment of epitope-associated nonsynonymous mutations among strongly selected mutations as compared to random chance. The fold enrichment values obtained via LDEN are also higher compared to LD-agnostic, which ignores the LD between mutations. The solid lines demonstrate the actual values of the fold enrichment, whereas the shading represents the standard deviation obtained using non-parametric bootstrap resampling.

## 4 Discussion

In this work, we have presented LDEN, a novel method for direct estimation of LD from NGS data without relying on population reconstruction. We have demonstrated that LDEN performs well under conditions of high read coverage and read length, which are the primary determinants of observable LD information. We have also demonstrated that LDEN substantially outperforms population reconstruction-based LD estimation both in terms of accuracy of the estimated LD values as well as the wall clock time, achieving a speedup of two orders of magnitude. Finally, we analyzed longitudinal norovirus data from an immunocompromised patient with LDEN and demonstrated that the inferred insights are biologically meaningful.

LDEN performs well for high coverage and long read length, which can be understood by the underlying parameters that change with these. High coverage reduces the statistical noise in the observed LD values. This statistical noise scales as 1*/n*_*i j*_, and increasing coverage reduces this noise, which in turn, reduces the error in the observed LD values. Longer read lengths, conversely, increase the range of observable LD entries that satisfy *j* − *i < r*, which renders more of the LD matrix directly observable rather than in need of estimation.

The superior performance of LDEN compared to MPL-R is attributed to fundamental differences in problem formulation.

MPL-R solves a harder problem than is necessary, that is, it first reconstructs complete sequences and then obtains LD from these. This approach is computationally expensive. Population reconstruction is also sensitive to low coverage and introduces errors that propagate into the LD estimates. LDEN, by contrast, directly computes the required two-point statistics from the reads and formulates the task as a convex matrix completion problem. This change of formulation is advantageous because it is computationally tractable, imposes only those constraints that are mathematically required for the task, and is guaranteed to converge to the unique global minimum because of the intersection of two convex sets. This fundamentally different and more efficient approach leads to the observed two orders of magnitude speedup. LDEN also remains scalable with increasing genome length (Supplementary Fig. S4). However, the performance degrades as the genome length is increased for the same read length because the amount of unobservable LD increases (Supplementary Section S1.3).

LDEN also provides a biologically meaningful rank ordering of the selection coefficients in the norovirus VP1. The statistically significant fold enrichment in epitope-associated nonsynonymous mutations identified by LDEN provides biological validation of the method.

While LDEN addresses some limitations of existing approaches, some challenges remain. APM, which forms the base for LDEN, is known to become slow to converge as the scale increases. This is because optimizing for one constraint may violate the other and vice versa, leading to a “zig-zagging” behavior which takes longer to stabilize for larger systems. The convergence can be accelerated via the incorporation of Dykstra’s correction^25^, which maintains a memory term that stores the difference between the input matrix and its projection onto the positive semidefinite constraint. At each iteration, this memory term is subtracted from the current matrix before the next projection, effectively correcting for overshoot and helping reach convergence faster. While the current implementation of LDEN works well for systems having up to 1500 bp genome length, future work can focus on implementing Dykstra’s correction to attain even greater speedup, particularly for large genomes sampled by longer reads for which the current implementation may become slow.

## 5 Acknowledgments

This work was carried out on the High Performance Computing Cluster provided by the Department of Electrical Engineering and CityU HK Burgundy High Performance Computing system managed and provided by the Computing Services Centre at City University of Hong Kong. The authors would like to express gratitude to Prof. Runsheng Li, Prof. Ka Chun Wong, Prof. Xiangjun Du, Dr. Muhammad Ashraf Abdelhamid Abdelgawad, and Dr. Abdurrashid Ibrahim Sanka for their insightful comments that improved the content of this study.

## 6 Conflict of interest

None declared.

## 7 Funding

This work was supported by the Hong Kong Innovation and Technology Fund (ITF) (MRP/071/20X), Applied Research Grant 9667256, and the Institute of Digital Medicine, City University of Hong Kong.

## 8 Data availability

The norovirus infection data can be accessed from the repository mentioned in the respective publication. Simulated data for testing LDEN can be accessed on the GitHub repository.

## 9 Software availability

The code for LDEN is available with example data on GitHub at https://github.com/SMUAbdullah/LDEN.

## Notes

### Competing Interest Statement

The authors have declared no competing interest.

