## supplementary_information for "LDEN: Alternating projections for whole genome linkage disequilibrium estimation from next-generation sequencing data"

**Table S1.** Comparison between the APM framework and LDEN.

| Parameter | APM | LDEN |
|---|---|---|
| Input | Partial correlation matrix | Partial covariance (LD) matrix |
| First constraint | Unit diagonal | Variance values in diagonal, known intra-read LD off-diagonals |
| Second constraint | Positive semidefiniteness | Positive semidefiniteness |
| Output | Completed correlation matrix | Completed covariance (LD) matrix |

**Table S2.** Prevalence values of the four variants used in datasets 1 to 4. $U(a,b)$ represents a uniform distribution over the interval $[a,b]$. "Rest" indicates the remaining prevalence after assigning prevalence values to all other variants.

| Dataset | Founder 1 | Founder 2 | Founder 3 | Founder 4 |
|---|---|---|---|---|
| 1 | $U(0.80,0.90)$ | $U(0.01,0.05)$ | Rest/2 | Rest/2 |
| 2 | $U(0.40,0.45)$ | $U(0.40,0.45)$ | Rest/2 | Rest/2 |
| 3 | $U(0.25,0.30)$ | $U(0.25,0.30)$ | $U(0.25,0.30)$ | Rest |
| 4 | $U(0.20,0.25)$ | $U(0.20,0.25)$ | $U(0.20,0.25)$ | Rest |

### S1 Supplementary results

#### S1.1 The error correction scheme improves the accuracy of the intra-read LD values

We tested whether our proposed scheme correctly suppresses sequencing errors and mitigates the effects of non-uniform coverage when calculating intra-read LD from NGS reads. Supplementary Figure S1 presents a boxplot of the Pearson correlation coefficient between the intra-read LD calculated from the reads using our proposed scheme, and the actual LD calculated for the same locus pairs from the full-length reference sequences. We first calculated the LD from the reads without either of the 1% frequency threshold and the shrinkage factor. These results are labeled "Unfiltered". We then applied the 1% frequency threshold and observed a significant improvement in accuracy. Then the shrinkage factor $\lambda$ was introduced and varied from 1 to 15. It can be observed that the proposed scheme with the frequency threshold and shrinkage helps improve the accuracy of the LD calculated from the reads, with consistently high correlation with the reference across all Monte Carlo runs. It can also be observed that the exact value of $\lambda$ is not important and any value about 5 gives similar results. Based on these results, we used $\lambda = 5$ in all our tests.

#### S1.2 LDEN demonstrates fast convergence

We set the convergence threshold $\varepsilon = 10^{-8}$ and chose the maximum number of iterations M from 100,1000,10000,100000. For M=100,1000, the convergence threshold was never reached and iteration always stopped when the number of iterations reached M, whereas for M=10000,100000, some Monte Carlo runs reached the convergence threshold (Supplementary Fig. S2A). The Frobenius distance between the last and the previous iterate decreased rapidly with an increase in M (Fig. S2B), however in terms of correlation with the ground truth LD matrix, the performance saturated for M=1000 (Fig. S2C). The wall clock time increased as M was increased, however it never exceeded a few tens of seconds even for M=100000 (Fig. S2D). Overall, it can be observed that LDEN rapidly converges to LD matrices that are strongly correlated with the reference LD matrix, and increasing the number of iterations does not improve the results much. Based on these results, we set M=1000 in all subsequent tests.

#### S1.3 LDEN is scalable with increasing genome length

We evaluated the scalability of LDEN by comparing the wall clock time for different genome lengths. LDEN required a median wall clock time of 6.83 second to process genomes of length 1500 bp, the largest genome length tested (Supplementary Fig. S4). This was significantly faster than the wall clock time of MPL-R, which required more than tens of minutes to process genomes of only 500 bp. We did not test LDEN on longer genomes because the read length of 150 bp used in the tests is too short relative to larger genome lengths, which renders most of the LD information unobservable. Even for a genome length of 1500 bp, the median Pearson correlation coefficient of the recovered LD was only slightly above 0.5, indicating that the limiting factor in scalability is not the wall clock time of the method but rather the amount LD information that can be observed by individual reads.

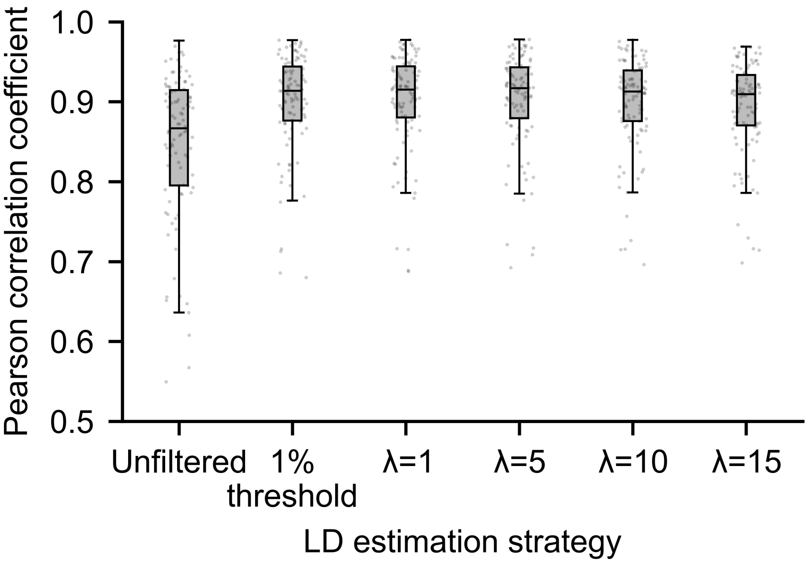


**Figure S1.** Proposed error correction scheme yields accurate intra-read LD. Boxplots of the Pearson correlation coefficient between the intra-read LD values obtained directly from the NGS reads and the LD values obtained from the full-length sequences demonstrate that LD values estimated using the proposed correction scheme are more accurate compared to those obtained without the correction. "Unfiltered" refers to no correction. "1% threshold" refers to filtering out loci with less than 1% variation in the single mutant allele frequencies. The rest of the plots show the results with the variation filtering as well as linear shrinkage. The results are shown for 100 Monte Carlo runs. Each Monte Carlo run consisted of a population of 1000 sequences derived from 4 variants. One founder of length 500 bp sequence was generated randomly while 3 variants were generated from the founder sequence by randomly mutating 1%, 2%, and 5% of the loci, respectively. Prevalence values of each variant follow dataset 1 in Supplementary Table S2. Analysis was performed on single-end NGS reads of 150 bp length with a coverage of 1500X.

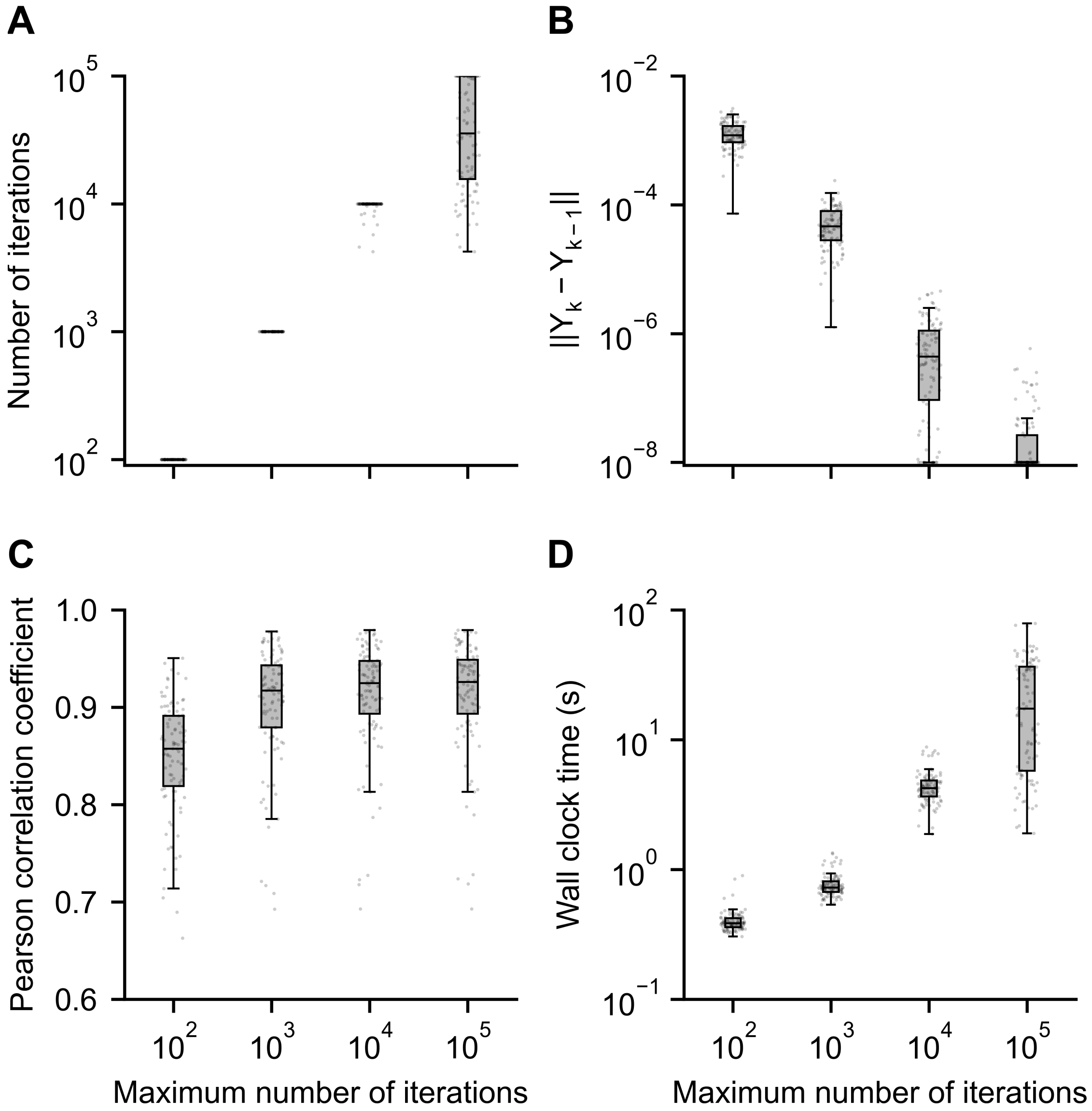


**Figure S2.** LDEN demonstrates fast convergence. **(A)** The maximum number of iterations is reached before the Frobenius distance between the last two iterates reaches the threshold of $10^{-8}$ for low values of the maximum number of iterations. **(B)** The Frobenius distance between the last two iterates decreases as the maximum number of iterations is increased. **(C)** The Pearson correlation coefficient between the estimated LD matrices and the reference LD matrices does not improve much with the increase in the number of iterations after reaching a peak value. **(D)** The wall clock time increases as the maximum number of iterations is increased but does not exceed a few minutes. Results are shown for 100 Monte Carlo runs. Each Monte Carlo run consisted of a population of 1000 sequences derived from 4 variants.One founder of length 500 bp sequence was generated randomly while 3 variants were generated from the founder sequence by randomly mutating 1%, 2%, and 5% of the loci,respectively. Prevalence values of each variant follow dataset 1 in Supplementary Table S2. Analysis was performed on single-end NGS reads of 150 bp length with a coverage of 1500X.

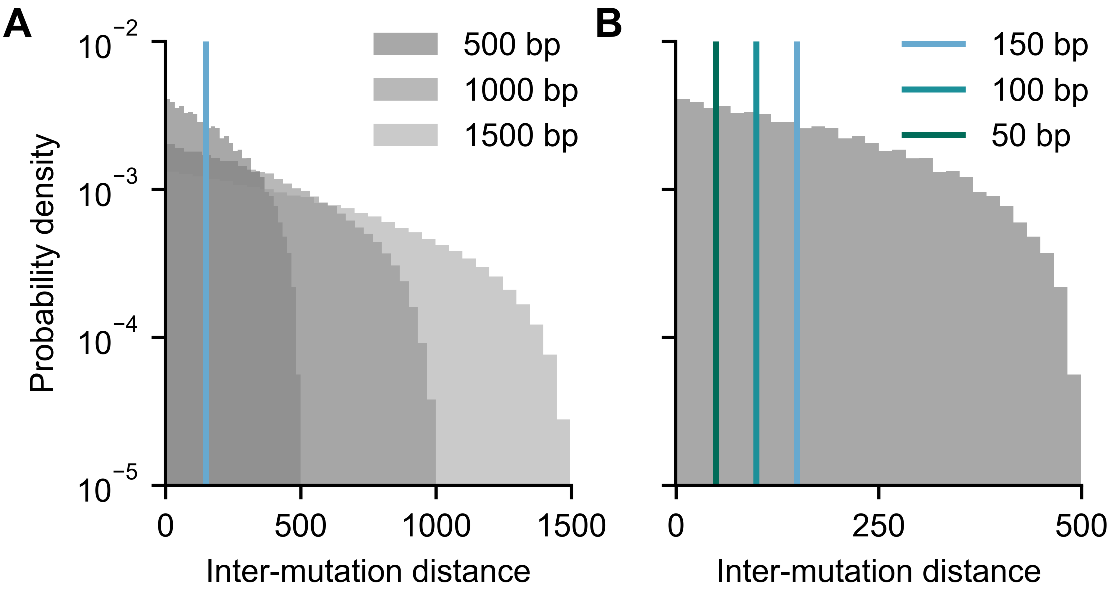


**Figure S3.** The information observable via reads depends on the read length and the genome length. **(A)** For a fixed read length of 150 bp (vertical blue line), increasing the genome length increases the number of unobservable LD values, as inter-mutation distances grow larger relative to read length. LD values corresponding to loci pairs to the left of the vertical line are observable via individual reads, whereas LD values to the right are unobservable. **(B)** When read length is increased while the genome length is held constant, more LD values become observable. The vertical lines represent the read length. LD values to the left of each line are observable via individual reads, whereas LD values to the right are unobservable. Results are shown for 100 Monte Carlo runs. Each Monte Carlo run consisted of a population of 1000 sequences derived from 4 variants. One founder sequence was generated randomly while 3 variants were generated from the founder sequence by randomly mutating 1%, 2%,and 5% of the loci, respectively. Prevalence values of each variant are listed in Supplementary Table S2. Genome lengths of the sequences were 500, 1000, 1500. Analyses were performed on simulated single-end NGS reads of 150 bp length with a coverage of 1500X.

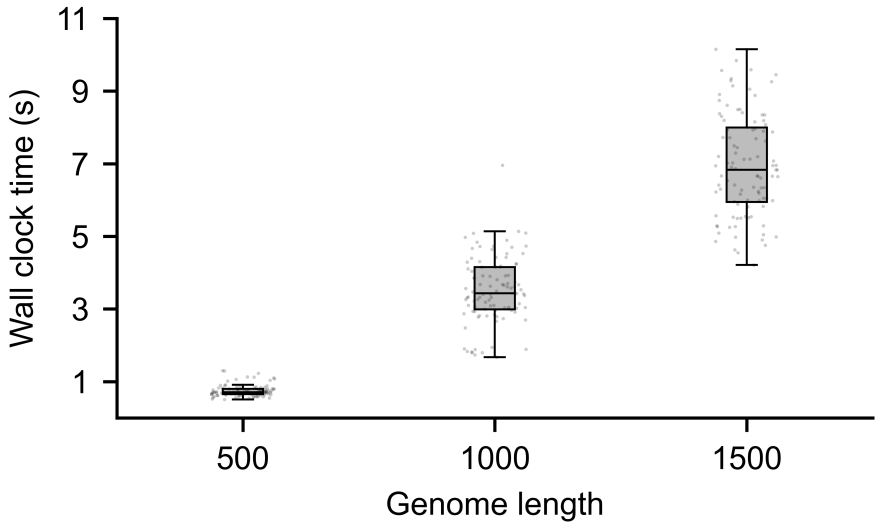


**Figure S4.** LDEN is scalable with increasing genome length. The wall clock time of LDEN increases with the genome length, but it does not increase significantly. Results are shown for 100 Monte Carlo runs. Each Monte Carlo run consisted of a population of 1000 sequences derived from 4 variants. One founder sequence was generated randomly while 3 variants were generated from the founder sequence by randomly mutating 1%, 2%, and 5% of the loci, respectively. Prevalence values of each variant are listed in Supplementary Table S2. Genome lengths of the sequences were 500, 1000, 1500. Analyses were performed on simulated single-end NGS reads of 150 bp length with a coverage of 1500X.
